# A Live-Cell Assay for the Detection of pre-microRNA-Protein Interactions

**DOI:** 10.1101/2020.06.23.167734

**Authors:** Sydney L. Rosenblum, Daniel A. Lorenz, Amanda L. Garner

## Abstract

Recent efforts in genome-wide sequencing and proteomics have revealed the fundamental roles that RNA-binding proteins (RBPs) play in the life cycle and function of both coding and non-coding RNAs. While these methodologies provide a systems-level view of the networking of RNA and proteins, approaches to enable the cellular validation of discovered interactions are lacking. Leveraging the power of bioorthogonal chemistry- and split-luciferase-based assay technologies, we have devised a conceptually new assay for the live-cell detection of RNA-protein interactions (RPIs), RNA interaction with Protein-mediated Complementation Assay, or RiPCA. As proof-of-concept, we have utilized the interaction of the pre-microRNA, pre-let-7, with its binding partner, Lin28. Using this system, we have demonstrated the selective detection of the pre-let-7-Lin28 RPI in both the cytoplasm and nucleus. Furthermore, we determined this technology can be used to discern relative affinities for specific sequences as well as of individual RNA binding domains. Thus, RiPCA has the potential to serve as a useful tool in supporting the investigation of cellular RPIs.

## Introduction

Recent studies have shown that RNAs are invariably bound to and often modified by RNA-binding proteins (RBPs).^1–6^ Thus, it is no surprise that RBPs have been found to play key roles in regulating many aspects of coding and non-coding RNA biology, including RNA processing, nuclear export, cellular transport, function, localization, and stability.^1, 3, 7, 8^ These efforts are carried out by >1,500 unique RBPs that utilize a variety of RNA-binding domains (RBDs) to achieve oftentimes specific and high affinity interactions with target transcripts.^1, 2, 7–9^ Accordingly, disruption of this complex network of RNA-protein interactions (RPIs) has been implicated in a number of human diseases including neurodegenerative, neuromuscular, neurodevelopmental and cardiovascular diseases, encephalo-, ribosomo- and neuropathies, autoimmune disorders, and many types of cancer.^7, 10–13^ Thus, the targeting of RBPs and RPIs has arisen as a new frontier in RNA-targeted drug discovery.^14^

Much of our knowledge regarding RPIs has been generated via large-scale sequencing and proteomics efforts.^15–17^ Methods such as RNA-immunoprecipitation followed by sequencing (RIP-seq) and crosslinking followed by immunoprecipitation and sequencing (CLIP-seq) have enabled the protein-centric identification of the RNA targets of a select RBP.^18, 19^ On the other hand, RNA-centric approaches, which typically utilize terminally-tagged RNA substrates for affinity enrichment,^20^ have employed quantitative mass spectrometry for the proteomic discovery of proteins bound to a specific RNA.^21–24^ While both of these methods have proved overlapping, yet non-redundant,^23^ each provides a systems biology view of the networking between RNA and protein, warranting the need for downstream technologies for validating discovered interactions.

Traditionally, the validation of RPIs has been carried out via *in vitro* biochemical and biophysical methodologies.^25–28^ Although useful, these techniques are low-throughput and quite laborious as they require the expression and purification of RBPs. Moreover, as these experiments are performed outside of cells, a high rate of false hits may be observed: false negatives due to the lack of required RBP binding partners or RBP post-translational modifications, and false positives due to formation of non-physiological RPIs in solution.^8, 27, 29, 30^ Early cellular assays of RPIs relied on yeast three-hybrid systems for the high-throughput screening of RNA sequences that bind to a specific RBP; yet, like the *in vitro* methods, this approach is performed outside of human cells and relies solely on the physical properties of the RNA and protein, not their natural biological activities.^31–33^ Mammalian cell-based assays have been reported and primarily rely on fluorescence imaging-based approaches for the cellular detection of RPIs.^34–39^ These methods are also not without drawbacks as tools for validating RPIs, particularly for interactions with poorly characterized RBPs, due to reliance on antibodies for RPI detection. Additionally, several of these techniques require that the RNA-of-interest is labeled with a protein-binding RNA affinity tag, such as the MS2 hairpin, to enable the design of multicomponent RNA-protein complementation assays to detect cellular RPIs via fluorescence and, more recently, chemiluminescence.^34, 40, 41^ While these methods have been shown to successfully detect interactions between proteins and long RNAs like mRNAs and lncRNAs, as well as enable the discovery of new RPIs^40^, the addition of a large hairpin RNA tag precludes it from use with small RNA species and could interfere with RPIs by altering local RNA structure and dynamics and affect RNA processing and localization.^42^

Over the past few years, the Garner laboratory has developed high-throughput screening technology for the discovery of small molecule modulators of RNA biology, including RPIs.^43–46^ Most recently, we have developed a click chemistry-mediated complementation assay, a homogeneous platform in which signal is dependent upon RPI-driven protein complementation of a split luciferase engineered from NanoLuc (NanoLuc Binary Technology; NanoBiT), allowing for catalytic signal amplification upon detection of full-length RPIs.^47^ As our previously developed assay platforms were biochemical in nature, to further advance our ability to study and manipulate RNA biology, we became interested in the development of a live-cell assay for RPIs. Herein, we report our efforts toward the development of a conceptually new approach for detecting RPIs in live cells.

## Results and discussion

### Development of a live-cell assay for detecting RPIs

Leveraging a related bioorthogonal chemistry-based strategy previously utilized by our group^47^, and making use of Promega’s HaloTag (HT)^48^ and NanoBiT^49^ systems, we devised an approach to directly monitor RPIs in living cells called RNA interaction with Protein-mediated Complementation Assay, or RiPCA (Fig. 1). In brief, cells stably expressing SmBiT-HT (step 1) are transiently transfected with an RBP-LgBiT plasmid and RNA probe which contains a PEGylated chloroalkane motif (step 2). In the cell, the RNA probe is first labeled with SmBiT through a covalent reaction between HT and the chloroalkane handle on the RNA (step 3). Successful interaction between the RNA and RBP is then monitored via chemiluminescence (step 5), which is produced upon SmBiT--LgBiT reassembly, forming the functional NanoLuc protein (step 4) (Fig. 1).

**Figure 1.**
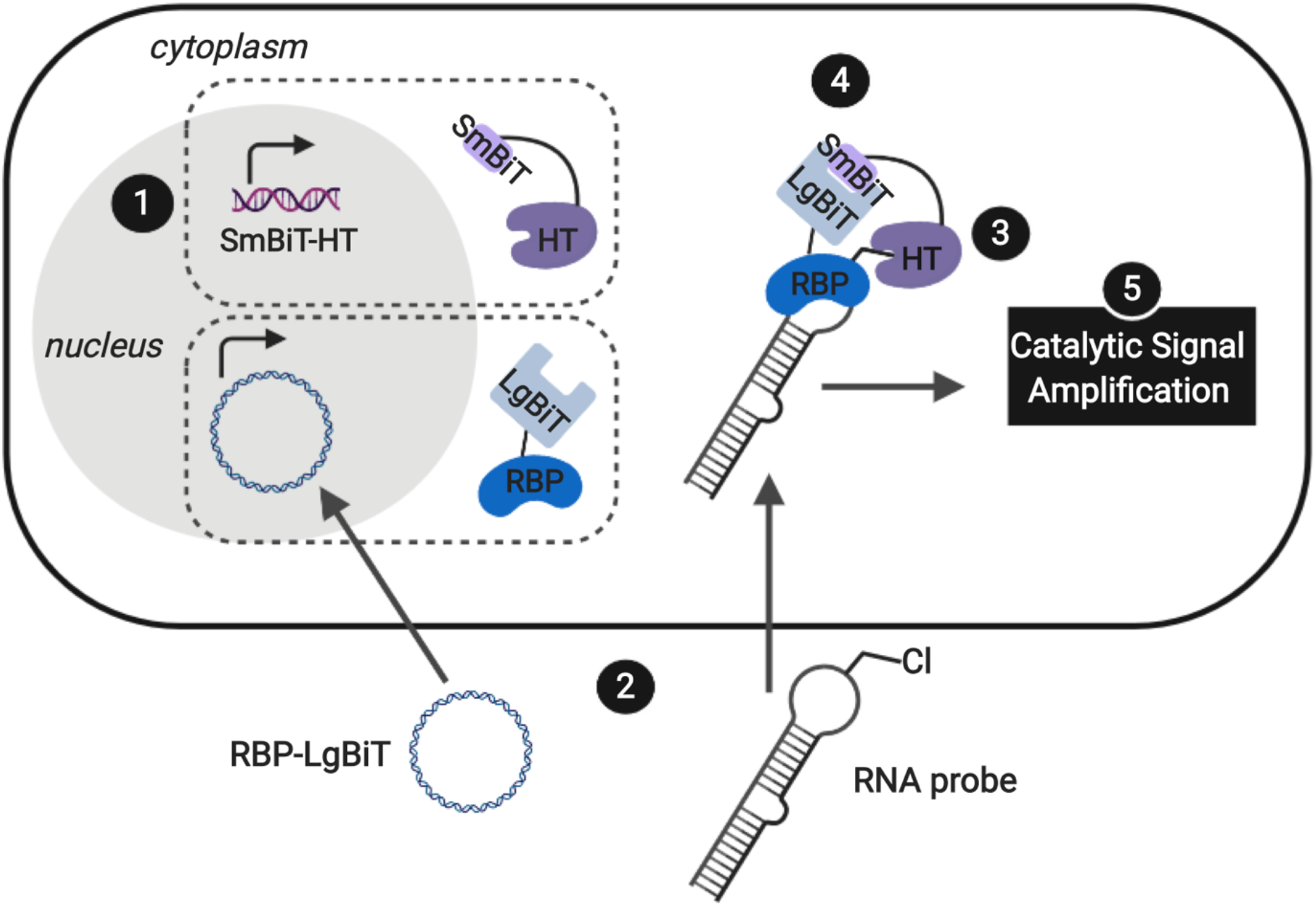
Live-cell assay for the detection of RPIs. In RiPCA, cells stably expressing SmBiT-HT (step 1) are transfected with an RBP-LgBiT plasmid and chloroalkane labelled-pre-miRNA probe (step 2). HT covalently labels the RNA probe with SmBiT (step 3) and upon interaction of the RBP and RNA-HT-SmBiT, the BiTs reassemble forming functional NanoLuc (step 4), which when treated with a luciferase substrate results in chemiluminescence (step 5). Created with BioRender.com.

RiPCA differs from the previously reported complementation assays for RPIs^41, 50–53^ in several important ways: (1) The weak affinity of SmBiT and LgBiT K_*d*_ of 190 μM) ^49^ ensures that signal generation is driven by the RPI, thereby enabling quantitative and accurate analysis. (2) Because our technology does not rely on the use of a protein-binding RNA affinity tag (e.g. MS2 hairpin) for RPI detection,^34, 40, 41, 54, 55^ we have the capability to detect RPIs beyond those involving mRNAs, including interactions with small non-coding RNAs such as miRNAs. (3) By using a NanoLuc-based format, enhanced sensitivity will be observed due to catalytic generation of chemiluminescence signal to promote favorable assay statistics. (4) The association of SmBiT and LgBiT is reversible demonstrating potential for the manipulation of the RPI using small molecule inhibitors or other cellular stimuli.^49^ Thus, we envisioned that the development of RiPCA would serve as a useful tool for the validation and study of cellular RPIs.

Based on our previous work in investigating the RPI between the pre-miRNA, pre-let-7, and RBP, Lin28,^44, 47^ we used this as a model for assay development. By binding to the terminal loop of primary or precursor forms of select let-7 family members, Lin28 inhibits maturation, and in some cases induces degradation, reducing levels of mature let-7.^56, 57^ Crystal structures of Lin28 in complex with the pre-element of pre-let-7 have been reported,^24, 57, 58^ which revealed that the protein uses three RNA-binding domains to interact with the hairpin loop of pre-let-7: an N-terminal cold shock domain (CSD) and two CCHC zinc knuckle domains (ZKD).^58^ The CSD and tandem ZKDs are connected by a flexible linker, which allows Lin28 to bind to the diverse let-7 family members, as well as its mRNA targets.^58^ To generate an RNA substrate, we used a modified pre-let-7d containing an aminoallyluridine substitution at U36 in the loop region that was subsequently converted into a PEGylated chloroalkane, herein referred to as pre-let-7d-Cl (Fig. S1).^47^ We hypothesized that this modification would be unlikely to disrupt interaction with Lin28 since it is outside of both the CSD and ZKD binding sites.^56–58^

To minimize the number of reagents that require transfection, a stable cell line expressing SmBiT-HT was generated from Flp-In HEK 293 cells. The 293-SmBiT-HT cells were then transiently transfected with pre-let-7d-Cl and LgBiT-tagged Lin28A constructs and grown in a 96-well plate. Both N- and C-terminally tagged Lin28A constructs were tested to determine if the location of the LgBiT tag would affect Lin28A binding to pre-let-7d-Cl. After incubation for 24 h, the media was changed, the cells were treated with NanoGlo Live Cell reagent and chemiluminescence resulting from the RPI was measured.

Excitingly, as shown in Fig. 2A, our proof-of-concept was successful and the assay selectively reported the interaction of pre-let-7d-Cl with both LgBiT-labeled Lin28A constructs over a LgBiT only control (Fig. 2A). We also demonstrated the necessity of the chloroalkane modification, as minimal signal was detected with the corresponding non-reactive RNA probe, which bears a trans-cyclooctene (TCO) handle in the terminal loop (Fig. 2A).^45^ Similar levels of chemiluminescence signal enhancement were observed with both Lin28A-LgBiT constructs, and we chose to use the C-terminally tagged protein in further experiments.

**Figure 2.**
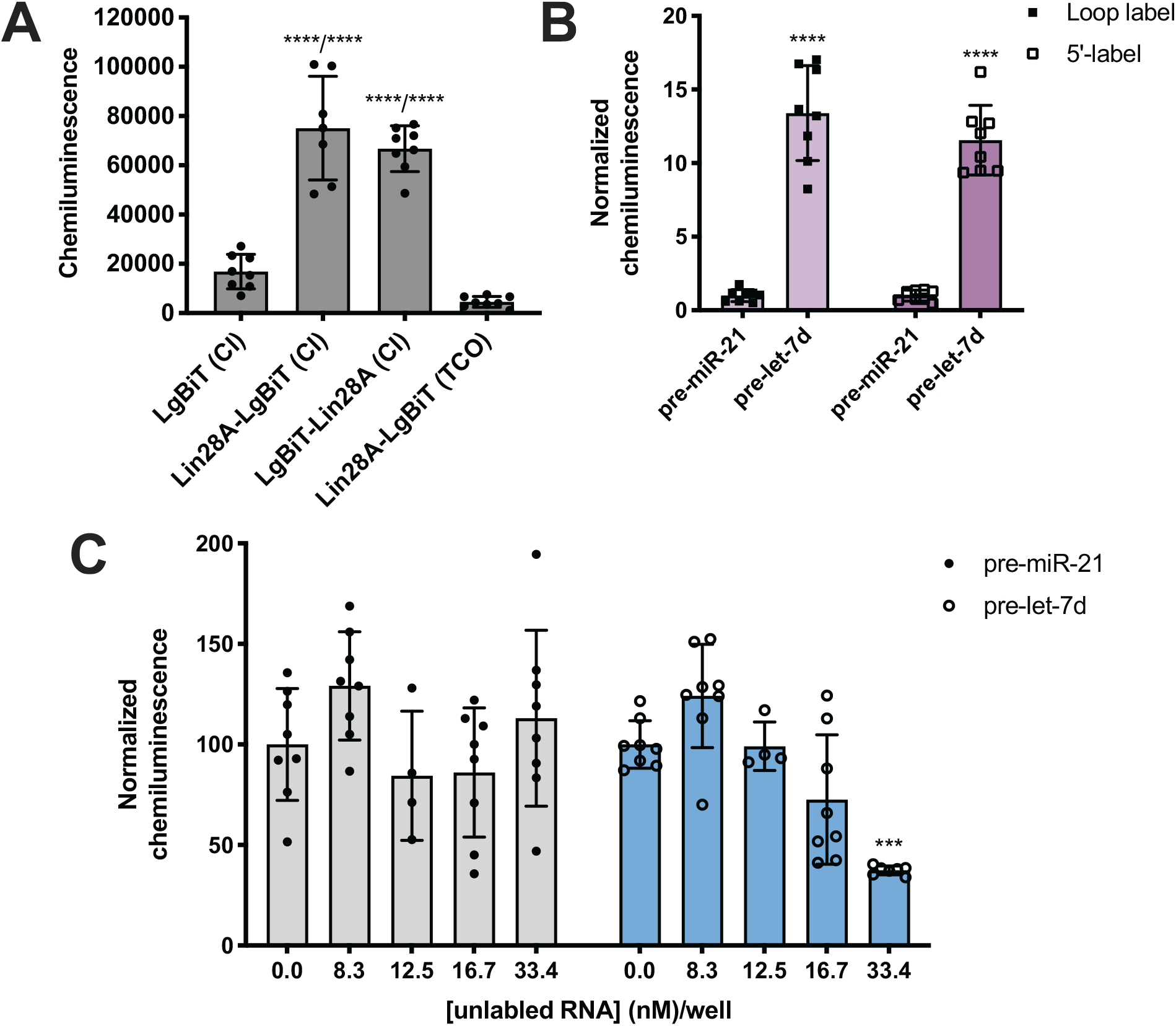
Proof-of-concept. (A) Selective detection of the Lin28A--pre-let-7d RPI using RiPCA. (B) The effect of varying the location of the chloroalkane linker. (C) Selective inhibition of pre-let-7d signal in RiPCA by titrating in increasing concentrations of an unlabeled pre-miRNA probe. See Tables S1-S3 for *p* values for each set of experiments.

We next turned our attention to the RNA substrate. To confirm that signal was dependent upon the selective interaction of Lin28A with pre-let-7d, we included a negative control sequence, pre-miR-21, that is not known to interact with Lin28A.^24^ Notably, signal produced by pre-miR-21 was 12—14-fold less than that of pre-let-7d, indicating that RiPCA can selectively detect the pre-let-7d—Lin28A RPI (Fig. 2B). As such, we used pre-miR-21 treatment as a measure of assay background in subsequent experiments. Wondering whether the location of the chloroalkane modification might impact signal detection, we generated additional pre-miRNA probes that placed the chloroalkane modification at the 5’ terminus of the hairpin. As shown in Fig. 2B, the location of the modification did not result in a significant difference in signal enhancement for pre-let-7d, nor a notable difference in raw signal (Fig. S3); negligible signal was again observed with pre-miR-21 demonstrating assay specificity.

With our ideal Lin28A and pre-let-7d substrates in hand, we examined the dynamic range of the assay by adjusting the amount of RBP-LgBiT plasmid and pre-let-7d-Cl transfected per well. Interestingly, as the amount of RBP-LgBiT plasmid transfected per well was increased, the signal-to-background (S/B) observed between pre-let-7d and pre-miR-21 decreased (Fig. S4A). This is likely a result of increased instances of non-specific interactions between SmBiT and LgBiT due to elevated cellular concentration of Lin28A-LgBiT. In contrast, increasing the amount of pre-miRNA-Cl transfected per well resulted in a concentration-dependent increase in S/B (Fig. S4B). From these results, and with the goal of maximizing S/B while minimizing the amount of transfection reagent used in each experiment, we proceeded with loop labeled pre-miRNA probes and transfected minimal quantities of the RNA and RBP reagents.

Finally, to demonstrate that RiPCA signal is produced as a result of ternary complex formation between pre-let-7d-Cl, SmBiT-HT, and Lin28A-LgBiT, cells were treated with increasing amounts of an unlabeled pre-miRNA, pre-let-7d or pre-miR-21. Importantly, we observed that unlabeled pre-let-7d, but not pre-miR-21, inhibited signal generated by treatment with the corresponding chloroalkane-labeled RNA probe (Fig. 2C). These data indicate that labeling of RNA with SmBiT does not contribute to non-specific interactions, further demonstrating that the specificity and strength of the signal in the assay is a direct measure of Lin28A binding to its target pre-miRNA. Encouraged by these promising data, we next set out to determine whether our read-out reflects Lin28 binding specificity by performing RiPCA with a variety of pre-miRNA probes (Fig. 3A). Again, we observed no signal in the presence of pre-miR-21, as well as pre-miR-34a, another sequence that has not been shown to interact with Lin28A (Fig. 3B).^24^ Additionally, we examined the interaction between Lin28A and other pre-let-7 isoforms, pre-let-7a-1 and pre-let-7g, which belong to different sub-classes of pre-let-7s. Analysis of let-7 sequences enriched in recent CLIP experiments revealed differential binding of distinct pre-let-7s by Lin28.^56^ This study found that Lin28 more strongly interacts with and regulates pre-let-7s that contain both a CSD and ZKD binding site (CSD^+^) compared to those that only contain a ZKD binding site (CSD^−^) (Fig. 3A).^56, 57^ This finding was corroborated in a proteomics study that identified RBPs enriched with specific pre-miRNA sequences, which demonstrated that Lin28 interacted primarily with CSD^+^ pre-let-7s.^24^ In congruence with these data, the two CSD^+^ sequences that were tested using RiPCA, pre-let-7d and pre-let-7g, produced robust chemiluminescence signal compared to pre-let-7a-1, a CSD^−^ sequence (Fig. 3B, Lin28A).^56^ Moreover, RiPCA measured greater signal generation with pre-let-7d in comparison to pre-let-7g (S/B of 13.4 and 6.2, respectively), which is correlative with interactome capture data wherein Lin28A was more highly enriched with pre-let-7d than pre-let-7g.^56^

**Figure 3.**
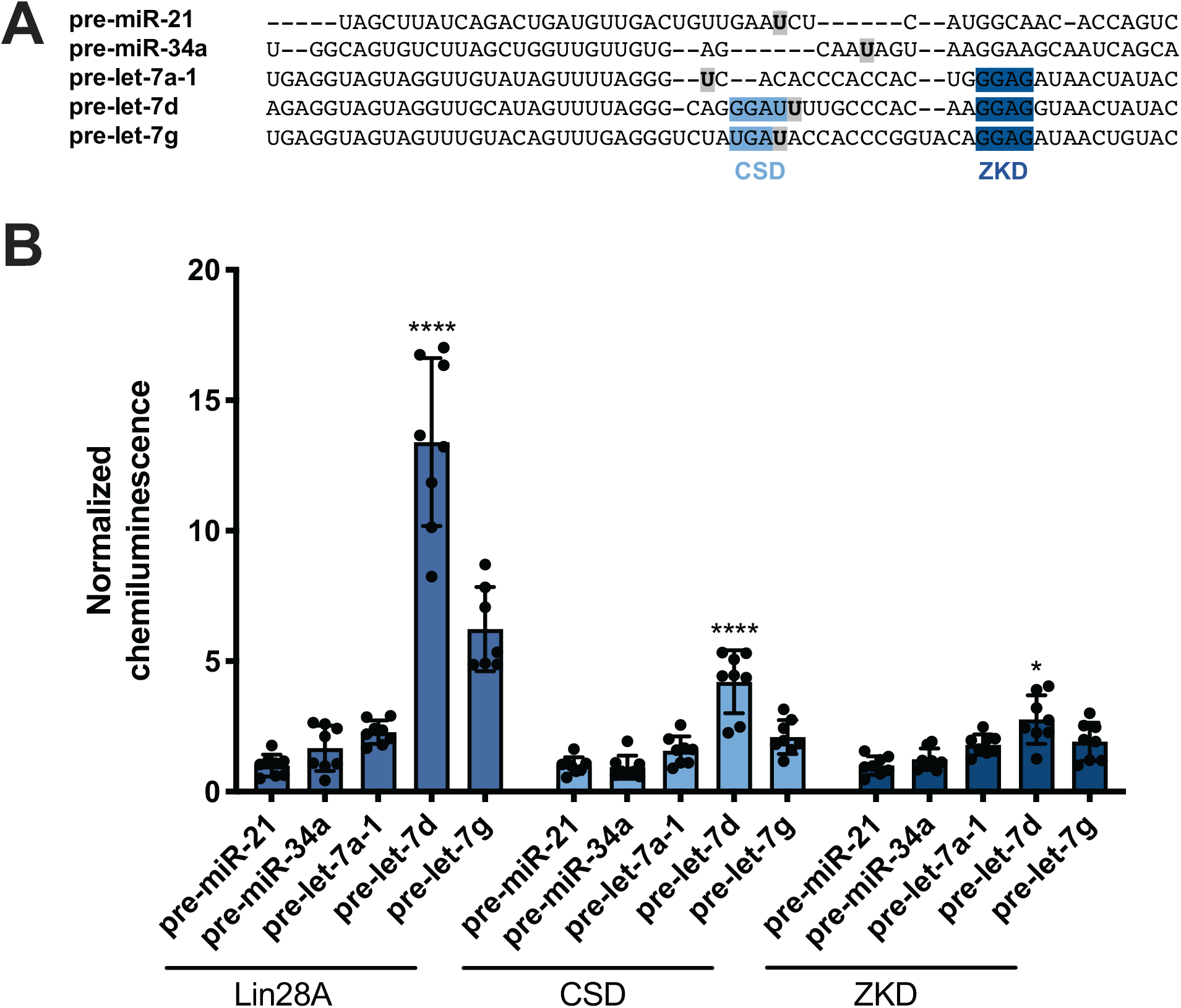
Selectivity of Lin28A in RiPCA. (A) Sequences of pre-miRNA probes used in RiPCA. The CSD and ZKD binding sites are highlighted in light and dark blue, respectively. The location of each aminoallyluridine modification is marked with a bold U highlighted in grey. (B) Selectivity of Lin28A-LgBiT, CSD-LgBiT, and ZKD-LgBiT in RiPCA. See Table S4 for *p* values for each set of experiments.

Since these results indicated that RiPCA is able to differentiate relative affinities of Lin28A binding to various pre-let-7 sequences, we next tested if RiPCA could similarly be capable of measuring the specificity of Lin28A’s RBDs and their relative affinities for pre-let-7s. Previous characterization of the CSD and ZKD have suggested that, while the ZKD may show higher sequence specificity, the CSD has much higher affinity for pre-let-7s and is a major determinant in the specificity of binding of full-length Lin28A.^57, 58^ Excitingly, in RiPCA, we observed that the relative binding preferences of the CSD for the various pre-miRNA sequences was similar to that of full-length Lin28A, whereas there was little difference in detected interaction of the ZKD with any of the various pre-miRNA sequences (Fig. 3B). Based on *in vitro* binding studies, Lin28A was found to bind to pre-let-7g with a K_*d*_ of ~50 nM, whereas the CSD and ZKD affinities were ~4- and ~15-fold weaker than Lin28A with K_*d*_s of ~195 nM and ~750 nM, respectively.^57, 58^ Notably, in RiPCA, the CSD produced much lower S/B for pre-let-7g, as well as pre-let-7d (S/B of 2.8 and 4.2, respectively), which is a ~3—5-fold reduction in comparison to Lin28A, matching the relative binding affinities measured *in vitro* (Fig. 3B).^57, 58^ Also, in line with the hypothesis that RiPCA can differentiate between weak and tight RPIs, the signal produced by the ZKD in RiPCA was lower than that of the CSD; however, the fold reduction in S/B produced by the ZKD compared to Lin28A was much smaller (3.0—3.3-fold) than *in vitro* measurements (~15-fold) (Fig. 3B). It is possible that the affinity threshold of the assay is lower than that of the ZKD for its let-7s, which presents an affinity limitation of the technology. Another possibility is that the ZKD is not able to compete with endogenously expressed Lin28B for binding to pre-let-7s in Flp-In HEK293 cells.

While Lin28A is not endogenously expressed in Flp-In HEK293 cells, these cells do express the closely related homologue, Lin28B. Lin28B is slightly larger than Lin28A; however, the CSD and ZKD remain highly conserved and are reported to interact with the same RNA sequences.^56^ Thus, RiPCA was applied to the detection of the interaction between Lin28B-LgBiT and the various pre-miRNAs. Consistent with the Lin28A RiPCA data, Lin28B-LgBiT interacted with CSD^+^, but not CSD^−^ pre-let-7s, and we again observed higher signal with pre-let-7d compared to pre-let-7g (S/B of 12.3 and 8.4, respectively) (Fig. 4). From these results, we were encouraged that RiPCA is not inhibited by the presence of an endogenous RBP, which opens up the opportunity to utilize RiPCA to detect a wide range of cellular RPIs.

**Figure 4.**
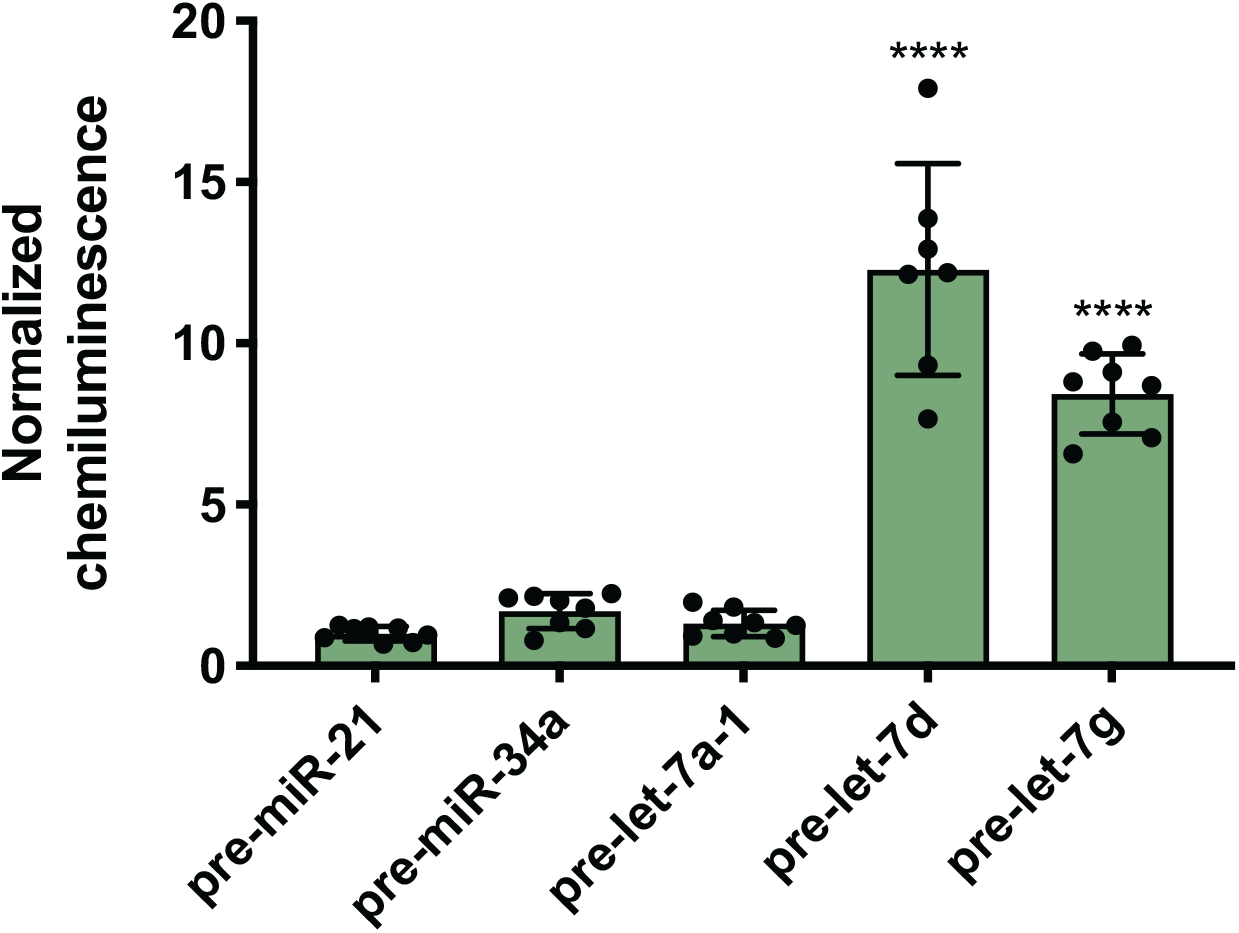
RiPCA with Lin28B. Selectivity of Lin28B-LgBiT against pre-miRNAs in the cytoplasm. See Table S5 for *p* values.

### Nuclear RiPCA

Next, we were excited to explore the possibility of detecting RPIs in distinct cellular compartments. As many RBPs function, at least in part, within the nucleus, we set out to develop a nuclear RiPCA detection system (nuc-RiPCA). A SmBiT-HT construct fused to the nuclear localization sequence (NLS) of SV40 large T antigen at the C-terminus was designed and stably expressed within Flp-In HEK293 cells. As confirmation, we performed Western blot and confocal microscopy, which revealed expression and cytoplasmic and nuclear localization of SmBiT-HT and SmBiT-HT-NLS, respectively, upon treatment with a tetramethylrhodamine (TMR)-labeled HT ligand and a nuclear stain (Fig. 5 and Fig. S5).

**Figure 5.**
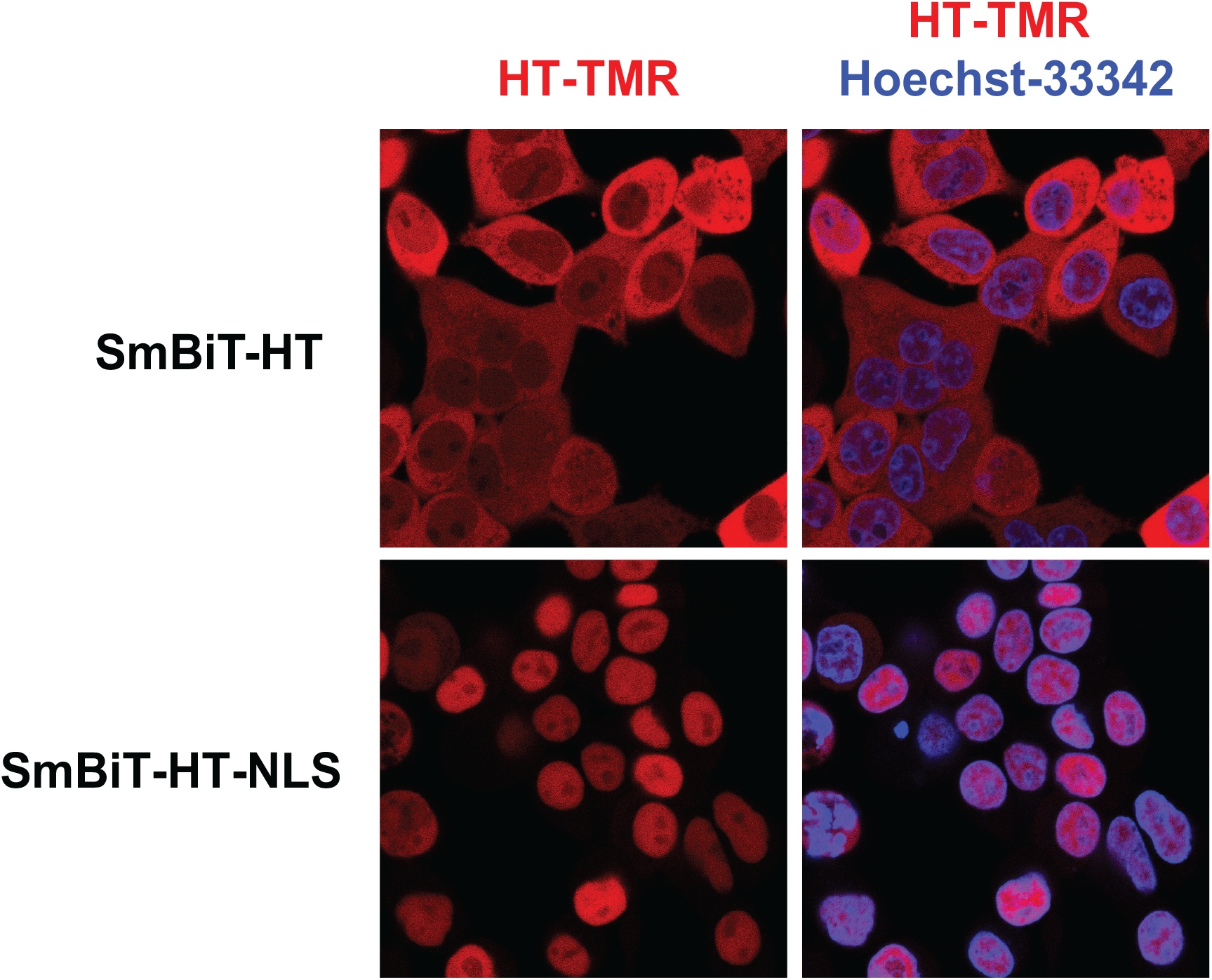
Localization of SmBiT-HT. Detection of SmBiT-HT and SmBiT-HT-NLS in Flp-In HEK293 cells via confocal microscopy. SmBiTs were detected following conjugation with a tetramethylrhodamine (TMR)-labelled HT ligand (red). Nuclei were stained with Hoechst 33342 (blue). Images on the left are of the SmBiT-HTs alone; those on the right are overlaid with the nuclear stain.

Because Lin28A and Lin28B have been shown to function as let-7 repressors both within the cytoplasm and nucleus,^59, 60^ we again used this as a model. Using the SmBiT-HT-NLS expressing cell line, we detected the production of chemiluminescence signal indicative of the interaction of Lin28A-LgBiT, as well as LgBiT-Lin28A, with pre-let-7d (Fig. 6 and Fig. S6). Relative to cytoplasmic RiPCA, we observed a 2.7- and 2.2-fold reduction in raw chemiluminescence signal produced upon interaction of pre-let-7d with Lin28A-LgBiT and LgBiT-Lin28A, respectively (Fig. S6).

**Figure 6.**
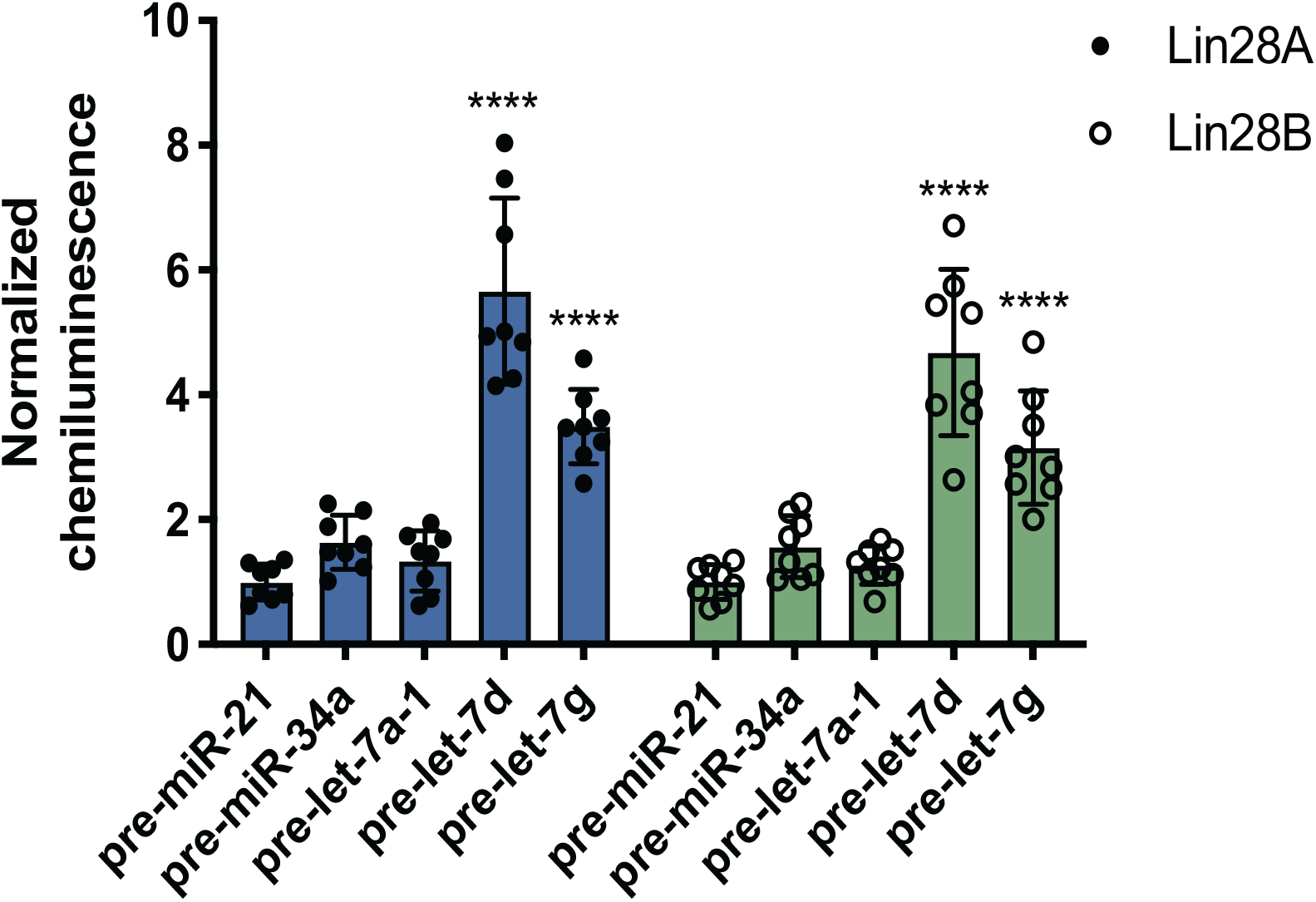
Localization of SmBiT-HT. Selectivity of Lin28A-LgBiT (left) and Lin28B-LgBiT (right) against various pre-miRNAs. See Table S6 for *p* values.

We were then interested in exploring whether we could tune the signal in nuc-RiPCA by adjusting the amount of DNA and pre-miRNA-Cl transfected. In contrast to cytoplasmic RiPCA, increasing the amount of DNA transfected resulted in greater signal relative to the pre-miR-21 control (Fig. S7A). With respect to RNA transfection, nuc-RiPCA yielded the same trend as in the cytoplasm, and S/B increased with increasing pre-miRNA-Cl (Fig. S7B).

Following this successful result, we tested nuc-RiPCA against other pre-miRNAs with both Lin28A-LgBiT and Lin28B-LgBiT. In accordance with our results obtained using the cytoplasmic assay, we observed signal only with the two CSD^+^ sequences, pre-let-7d and pre-let-7g, with both Lin28A- and Lin28B-LgBiT (Fig. 6). In addition to observing lower overall signal in the nucleus than in the cytoplasm, we also found that CSD^+^ sequences produced S/B that was between 1.8- and 2.7-fold lower than in the cytoplasm (Fig. 3 and Fig. 6). We hypothesize that the lower signal and S/B, as well as increased tolerance of LgBiT expression, could be the result of short residence time of the LgBiT-tagged protein in the nucleus, competition with endogenous Lin28B in the nucleus,^59, 60^ or decreased pre-miRNA probe uptake into the nucleus. With respect to the latter, this is unlikely to be problematic, as molecules <40 kDa have been shown to readily traverse through the nuclear pore complex.^61^ We anticipate that the ability of RiPCA to enable organelle-specific RPI detection will open the door to its future application in the study of disease-relevant nuclear RBPs critical in human health.

## Conclusions

RBPs and RPIs play important, but currently understudied roles in the maintenance of human health. While the advent of sequencing and quantitative mass spectrometry has dramatically enhanced our ability to globally profile these interactions, revealing intricate and complicated networks of RPIs, experimental validation of these data sets remains a challenge. Using chemical biology- and bioorthogonal chemistry-based strategies, we have developed an innovative assay for the live-cell detection of RPIs, RiPCA. Through this approach, we have selectively detected the interaction of the Lin28 proteins and their pre-let-7 targets in both the cytoplasm and the nucleus. We also demonstrate that RiPCA is capable of discerning between high and low affinity RPIs, using the individual Lin28 RBDs as a model. Combined, these data provide encouraging proof-of-concept for this emerging technology. Efforts towards its adaptation to validate putative RBP--pre-miRNA interactions discovered via proteomics,^24^ as well as RPIs outside of those with pre-miRNAs, are currently being explored and will be reported in due course.

## Conflicts of interest

There are no conflicts to declare.

## Acknowledgements

This work was supported by the NIH (R01 GM118329 and R01 GM135252 to A.L.G.), the Dr. Ralph and Marian Falk Medical Research Trust (A.L.G.) and the University of Michigan Rackham Graduate School (S.L.R.).

## Notes

### Competing Interest Statement

The authors have declared no competing interest.

