## Supporting Information for "A Live-Cell Assay for the Detection of pre-microRNA-Protein Interactions"

**A. General Materials and Methods** Page S2

**B. Cloning** Page S2

**C. RNA Labelling** Page S4

**D. SmBiT-HT Stable Cell Lines** Page S5

**E. RiPCA Protocol** Page S5

**F. Confocal Microscopy** Page S6

**G. Supplemental Figures** Pages S7

**H. Supplemental Tables** Page S9

**I. References** Page S11

### A. General Materials and Methods

*General cell culture methods.* Flp-In™-293 cells stably expressing either SmBiT-HaloTag or SmBiT-HaloTag-NLS were grown in DMEM (Corning cat #10-017-CV) supplemented with 10% FBS (Atlanta Biologicals S11550), L-glutamine (Gibco cat #25030081), and hygromycin B (100 μg/mL) (Gibco cat #10687010) at 37 °C with 5% $CO_2$ in a humidified incubator, passaged at least once before use for an experiment. Cells were passaged using Trypsin-EDTA (0.25%) (Gibco cat #25300054) approximately 10 times, and no more than 15 times, before returning to low passage stocks. To count cells, cells were harvested and 10μL of the cell suspension was mixed with 10μL Trypan Blue (Gibco cat #15250061) ([final] = 0.2% trypan blue) and counted using a hemocytometer.

*General assay and data analysis methods.* Chemiluminescence data was collected on a BioTek Cytation3 plate reader. All data was analyzed using GraphPad Prism version 6.0c for Mac OS X (GraphPad Software, www.graphpad.com). All normalized chemiluminescence is reported as the signal of each well divided by the average signal of quadruplicate pre-miR-21 wells. The only exception is Fig. 2C, in which signal is normalized by dividing by the average of quadruplicate wells containing no unlabeled probe and multiplying by 100.

*Statistical analysis.* All statistical tests were performed using Prism (v8). One- or two-way ANOVA tests were run for each set of data. Details of multiple comparisons tests are included in table legends. Graphs show mean ± standard deviation.

*Materials.* Chemically synthesized pre-microRNAs (deprotected, desalted and HPLC purified), containing aminoallyl uridine or aminohexylacrylamino uridine modifications and biotin attached to the 5'-end of the sequence by an 18-atom spacer, were purchased from Dharmacon and used as received for the labeling reaction. HaloTag Succinimidyl Ester (O4) Ligand was purchased from Promega and used as received (cat #P6751). Note that the HaloTag Succinimidyl Ester (O4) Ligand should be dissolved and immediately portioned into single use aliquots stored at -80 °C to avoid degradation. Flp-In™-293 cells and associated vectors were purchased from ThermoFisher Scientific (Invitrogen cat #75007 and #601001, respectively). The Nano-Glo Live Cell Assay System was purchased from Promega and used as received (cat #N2012). HaloTag® TMR Ligand (Promega cat #G8251) and Lipofectamine™ RNAiMAX (Invitrogen cat #13778100) were used as received.

### B. Cloning

*SmBiT-HT cloning.* SmBiT-HT was amplified from a pFC30K construct containing a N-terminal SmBiT and C-terminal HaloTag (HT) fusion protein and inserted into pcDNA5/FRT using standard PCR cloning techniques with KpnI and NotI restriction enzymes.

Primers:

SmBiT-HT
5' GTACGGTACCGCCACCATGGTGACCGGCTACCGG
5' CAGTGCGGCCGCTCACTATTAGTGGTGATGGTGATGATG

*SmBiT-HT-NLS cloning.* To insert a C-terminal SV40 NLS, the multiple cloning site of the pcDNA5/FRT + SmBiT-HT construct was modified to include a Blp restriction site using standard PCR cloning techniques with KpnI and NotI restriction enzymes.

Primers:

SmBiT-HT-Blp
5’ GTACGGTACCGCCACCATGGTGACCGGCTACCGG
5’ GACTGCGGCCGCTCATTAGCTCAGCCCACCGGAAATCTCCAGAGTAG

The SV40 NLS (5’ CCAAAGAAAAAGAGAAAGTA) was then appended to the C-terminus of SmBiT-HT by annealing oligos and inserting them into pcDNA5/FRT + SmBiT-HT-Blp digested with Blp restriction enzyme.

Oligos:

SmBiT-HT-Blp
5’ TGAGTGGAGGTGGTCCAAAGAAAAAGAGAAAGTATGGC
5’ TCAGCCATACTTTTCTCTTTTTCTTTGGACCACCTCCAC

*Lin28A-LgBiT cloning.* Mouse Lin28A-LgBiT was amplified from pFN29K-Lin28A-LgBiT, previously cloned in the lab[1], and inserted into pcDNA3 using standard cloning techniques with KpnI and NotI restriction enzymes. Primers insert a Kozak sequence on the N-terminus.

Primers:

Lin28A-LgBiT
5’ GTACGGTACCGCCACCATGGCCTCGGTGTCCAACC
5’ CAGTGCGGCCGCTCATTAAACACTGTTGATGGTTACTCG

*LgBiT-Lin28A cloning.* First, a pcDNA3 construct with LgBiT at the N-terminal position was generated by amplifying LgBiT and insertion into pcDNA3 using standard cloning techniques with BamHI and XhoI restriction enzymes. Mouse Lin28A was amplified from the pcDNA3-Lin28a-LgBiT construct and ligated into a pcDNA3 construct with LgBiT inserted at the N-terminal position using standard cloning techniques with XhoI and XbaI restriction enzymes. Primers insert a Kozak sequence on the N-terminus.

Primers:

LgBiT
5’ GTAGGATCCGCCACCATGGTCTTCACACTC
5’ CATCTCGAGACTGTTGATGGTTACTC

Lin28A
5’ ATGCCTCGAGATGGGCTCGGTGTCCAACC
5’ CATCTAGAATTCTGGGCTTCTGGGAGCA

*Lin28B-LgBiT cloning.* Human Lin28B was amplified from pFC30K vector containing Lin28B and ligated into previously cloned pcDNA3 vector containing LgBiT inserted at the C-terminal position (see Lin28A-LgBiT cloning) using standard cloning techniques with KpnI and AsiSI restriction enzymes. Primers insert a Kozak sequence on the N-terminus.

Primers:

5’ ATGGTACCGCCACCATGGCCGAAGGCG
5’ ATGCGATCGCTGTCTTTTTCCT

### C. RNA Labelling

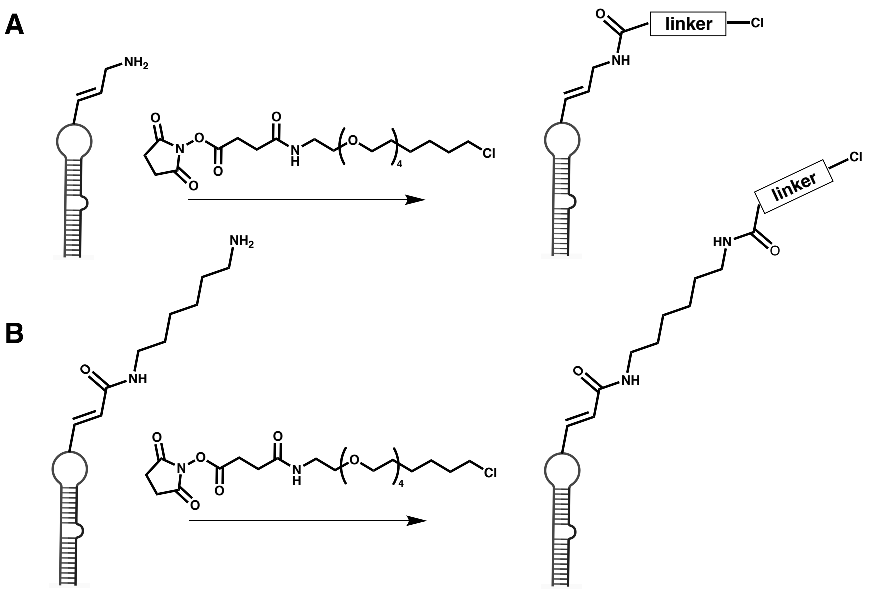


**Figure S1**. RNA labelling scheme with (A) 5-aminoallyl uridine and (B) 5-aminohexylacrylamino uridine modifications.

*Protocol.* Amino-modified pre-miRNA (1.0 mM in 100 mM phosphate buffer, pH 8.0) was mixed with an equivalent volume of HaloTag® Succinimidyl Ester (O4) Ligand (10 mM in DMSO). The reaction was then allowed to proceed at 25 °C for 1 h. pre-miRNA-Cl was precipitated by the addition of 0.11× volume of 3.0 M sodium acetate (pH 5.2) and 4 volume equivalents of cold ethanol, and pelleted at 20,000 × g for 40 min at 4 °C. The pellet was then re-suspended in 100 mM phosphate buffer (pH 8.0) at a concentration of 1.0 mM and stored at -80 °C.

For let-7s, the Lin28 CSD binding site is bolded and ZKD binding site is underlined.

**pre-miR-21 RNA Sequence:**
5'-Biotin-(18-atom spacer; hexaethylene glycol)-UAGCUUAUCAGACUGAUGUUGACUGUUG AA-(5-aminoallyl uridine)-CUCAUGGCAACACCAGUCGAUGGGCUGUC-3'

**pre-miR-34a RNA Sequence:**
5'-Biotin-(18-atom spacer; hexaethylene glycol)-UGGCAGUGUCUUAGCUGGUUGUUGUGAG CAA-(5-aminoallyl uridine)-AGUAAGGAAGCAAUCAGCAAGUAUACUGCCCUA-3'

**pre-let-7a-1 RNA Sequence:**
5'-Biotin-(18-atom spacer; hexaethylene glycol)-UGAGGUAGUAGGUUGUAUAGUUUUAGGG-(5-aminohexylacrylamino uridine)-CACACCCACCACUGGGAGAUAACUAUACAAUCUACUGU CUUUCU-3'

**pre-let-7d RNA Sequence:**
5'-Biotin-(18-atom spacer; hexaethylene glycol)-AGAGGUAGUAGGUUGCAUAGUUUUAGGG CAG**GGAU**-(5-aminoallyl uridine)-UUGCCCACAAGGAGGUAACUAUACGACCUGCUGCCU UUCU-3'

**pre-let-7g RNA Sequence:**
5'-Biotin-(18-atom spacer; hexaethylene glycol)-UGAGGUAGUAGUUUGUACAGUUUGAGGG UCUA**UGA-(5-aminohexylacrylamino uridine)-**ACCACCCGGUACAGGAGAUAACUGUACAG GCCACUG CCUUGCU-3'

**D. SmBit-HT Stable Cell Lines.** Flp-In-293 cells stably expressing a SmBiT-HT were generated by co-transfecting Flp-In-293 cells with 9 ng pOG44 and 1 ng pcDNA5/FRT using Lipofectamine LTX+ Plus reagent (Life Technologies) according to the manufacturer's instructions. Expression and localization of SmBiT-HaloTag or SmBiT-HaloTag-NLS were confirmed by Western blot and confocal microscopy.

**E. RiPCA Protocol**

*General protocol.* Flp-In-293 cells stably expressing a SmBiT-HT protein were reverse transfected using Lipofectamine™ RNAiMAX Transfection Reagent. Cells were passaged approximately 10 times, and no more than 15 times, before returning to low passage stocks. To test "n" number of conditions, Solution A was prepared by combining 50 × (n+1) µL of room temperature Opti-MEM and 2.4 × (n+1) µL plasmid encoding selected RBP-LgBiT fusion. Solution B was prepared by adding pre-miRNA-Cl and plasmid (final concentrations 0.3 µM and 0.195 ng/µL, respectively) to 50 µL Opti-MEM™ for each separate condition to be tested. Solution B was mixed with 50 µL of Solution A to yield Solution A+B, which was incubated for at least 15 min at room temperature while cells were harvested. Cells were harvested as and counted as described above. Harvested cells were used to prepare Solution C, which was composed of 400 µL of 100,000 cells/mL. Solution C was mixed with 50 µL of Solution A+B and plated 100 µL per well, four wells per condition, in a white-bottom, tissue culture-treated 96-well plate (Corning cat #3917). The plate was incubated in a humidified incubator (37 °C and 5% $CO_2$) for 24 h. After incubation, the media was removed and replaced with 100 µL room temperature Opti-MEM™ and treated with 25 µL NanoGlo Live Cell Reagent diluted 1:20 according to the manufacturer's recommendation. All chemiluminescence data was collected immediately on a BioTek Cytation3 plate reader.

Representative calculations based on an assay for n = 5 conditions:

Solution A: Prepared for n+1= 6
6 x 50 µL → 300 µL OptiMEM™
6 x 2.4 µL → 14.4 µL Lipofectamine™ RNAiMAX

Solution B:
50 µL OptiMEM™
2.5 µL 3.9 ng/µL RBP-LgBiT plasmid
0.3 µL 50 µM pre-miRNA-Cl

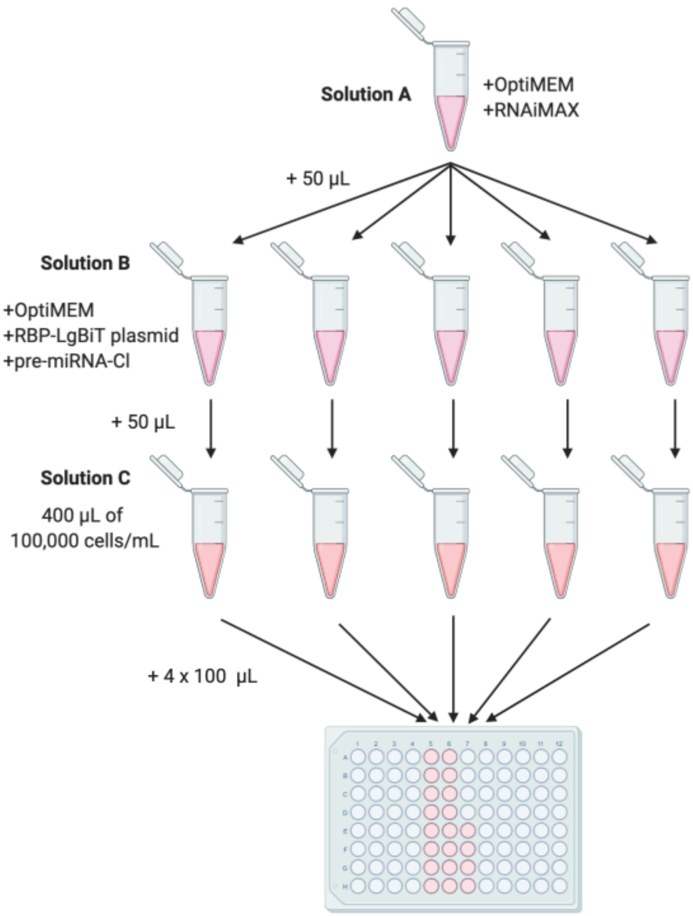


**Figure S2.** RiPCA transfection workflow.

*DNA and RNA titration.* To alter the amount of DNA transfected per well, higher concentration stocks of plasmid were used to allow addition of the same volume to each condition (final concentrations increased from 0.195 ng/µL to 0.39 ng/µL and 19.5 ng/µL). To alter the amount of RNA transfected per well, increasing volumes of 50 µM stock solution of pre-miRNA-Cl were used. The amount of Lipofectamine$^{TM}$ RNAiMAX was adjusted accordingly (from 2.4 × (n+1) µL for 16.7 nM/well to 1.2× (n+1) µL for 8.3 nM/well and 3.6 × (n+1) µL for 25 nM/well).

*Competition with unlabeled RNA.* In competition experiments, the general RiPCA protocol was followed with the following change. In addition to the DNA and RNA added to Solution B, varying amounts of unlabeled pre-miRNA were added to Solution B (0, 0.15, 0.225, 0.3, or 0.6 uL unlabeled probe).

### F. Confocal Microscopy

*Protocol.* Flp-In cells stably expressing SmBiT-HT or SmBiT-HT-NLS were harvested and counted using methods described above. Cells were diluted to a density of 100,000 cells/mL and 200 µL was plated in an 8-well chambered coverglass (Nunc$^{TM}$ Lab-Tek$^{TM}$ II). The chambered coverglass was incubated in a tissue culture incubator (37 °C and 5% $CO_2$) for 24 h to allow the cells to adhere to the glass. To stain live cells, the media was supplemented with a single stain or a combination of stains at final concentrations of 50 nM HaloTag® TMR Ligand (Promega cat #G8251), 0.44 μM Hoescht 33342 (Fisher), and 0.2 μM MitoTracker Green FM (Cell Signaling Technology). The chambered coverglass was returned to the incubator for 30 min. The media was then removed and replaced with 200 μL Opti-MEM™. Fluorescence was visualized using Nikon A1SI Confocal microscope. Images were processed with NIS-Elements.

### G. Supplemental Figures

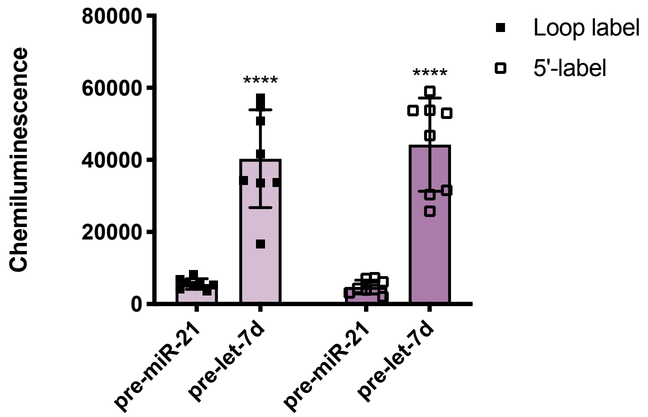


**Figure S3**. Chemiluminescence signal detection of interaction between Lin28A-LgBiT and 5'- or loop-labeled pre-miR-21 and pre-let-7d. See Table S2 for *p* values.

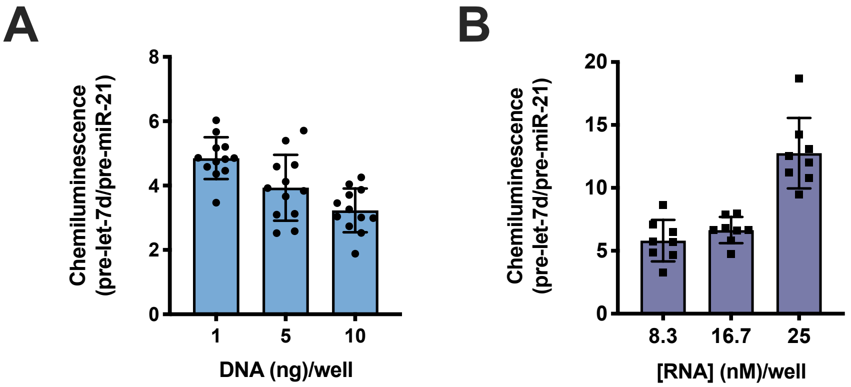


**Figure S4.** (A) Dependence of S/B on the amount of Lin28A-LgBiT plasmid and (B) pre-miRNA-Cl transfected in SmHT-expressing cells. Normalized chemiluminescence is reported as pre-let-7d/pre-miR-21.

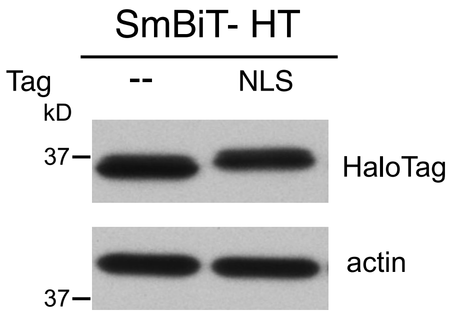


**Figure S5.** Expression of SmBiT-HT and SmBiT-HT-NLS constructs in stably expressing Flp-In 293 cell lines.

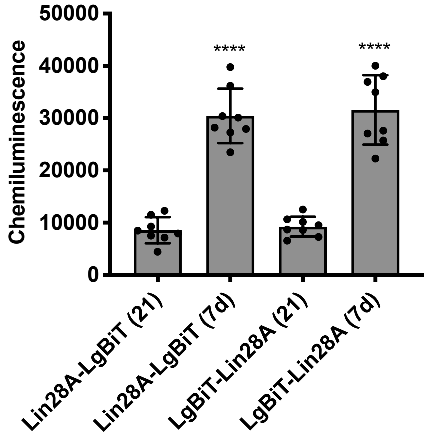


**Figure S6.** (A) Signal produced by SmHT-NLS-expressing cells co-transfected with pre-miR-21 or pre-let-7d and Lin28A tagged with LgBiT at the C- or N-terminus. Statistical significance determined by one-way ANOVA Sidak's multiple comparison test (n = 8); *p* <0.0001.

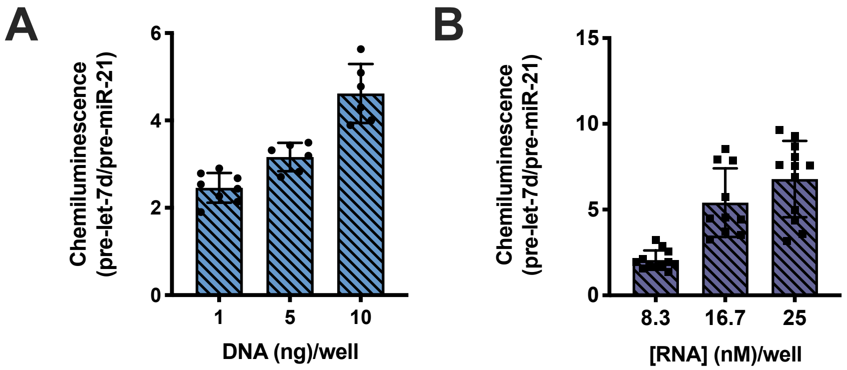


**Figure S7.** (A) Dependence of S/B on the amount of Lin28A-LgBiT plasmid and (B) pre-miRNA-Cl transfected in SmHT-cNLS-expressing cells. Normalized chemiluminescence is reported as pre-let-7d/pre-miR-21.

### H. Supplemental Tables

**Table S1.** Statistical significance associated with **Fig. 2A** between chemiluminescence produced by cells co-transfected with chloroalkane-labeled pre-let-7d and Lin28A-LgBiT or LgBiT-Lin28A and cells co-transfected with either a LgBiT or TCO-labeled pre-let-7d control (ns = not significant). Statistical significance determined by one-way ANOVA Sidak's multiple comparison test (n = 8).

| Construct | LgBiT control | TCO control |
|---|---|---|
| **Lin28A-LgBiT** | <0.0001 (****) | <0.0001 (****) |
| **LgBiT-Lin28A** | <0.0001 (****) | <0.0001 (****) |

**Table S2.** Statistical significance associated with **Fig. 2B and Fig. S3** between chemiluminescence produced by cells transfected with pre-miR-21 and pre-let-7d with chloroalkane labels in either the loop or at the 5'-end of the probe. Statistical significance determined by one-way ANOVA Sidak's multiple comparison test (n = 8).

| Construct | Loop label | 5'-label |
|---|---|---|
| **pre-let-7d** | <0.0001 (****) | <0.0001 (****) |

**Table S3.** Statistical significance associated with **Fig. 2C** between chemiluminescence produced by cells transfected with chloroalkane-labeled pre-miR-21 and pre-let-7d sequences and cells transfected with varying amounts of unlabeled pre-miR-21 and pre-let-7d in cytoplasmic RiPCA (ns = not significant). Statistical significance determined by one-way ANOVA Dunnett's multiple comparison test (n = 8, except 12.5, for which n = 4). One data point from the pre-let-7d [33.4] data set was identified as an outlier using the ROUT method where Q = 1% (Prism) and eliminated from the data set and statistical analysis.

| [Unlabeled RNA] (nM) | pre-miR-21 | pre-let-7d |
|---|---|---|
| **8.3** | 0.2845 (ns) | 0.5244 (ns) |
| **12.5** | 0.9738 (ns) | >0.9999 (ns) |
| **16.7** | 0.9554 (ns) | 0.3660 (ns) |
| **33.4** | 0.9690 (ns) | 0.0004 (***) |

**Table S4.** Statistical significance associated with **Fig. 3B** between signal-to-background (S/B) produced by various pre-miRNA-Cl sequences and pre-miR-21 with each LgBiT fusion in cytoplasmic RiPCA (ns = not significant). Statistical significance determined by two-way ANOVA Tukey's multiple comparison test (n = 8).

| Sequence | Lin28A | CSD | ZKD |
|---|---|---|---|
| **pre-miR-34a** | 0.7456 (ns) | >0.9999 (ns) | 0.9922 (ns) |
| **pre-let-7a-1** | 0.1464 (ns) | 0.8381 (ns) | 0.5962 (ns) |
| **pre-let-7d** | <0.0001 (****) | <0.0001 (****) | 0.0152 (*) |
| **pre-let-7g** | <0.0001 (****) | 0.2826 (ns) | 0.4645 (ns) |

**Table S5.** Statistical significance associated with **Fig. 4** between signal-to-background (S/B) produced by various pre-miRNA-Cl sequences and pre-miR-21 with Lin28B-LgBiT in cytoplasmic RiPCA (ns = not significant). Statistical significance determined by one-way ANOVA Sidak's multiple comparison test (n = 8).

| Sequence | Lin28B |
|---|---|
| **pre-miR-34a** | 0.8874 (ns) |
| **pre-let-7a-1** | 0.9929 (ns) |
| **pre-let-7d** | <0.0001 (****) |
| **pre-let-7g** | <0.0001 (****) |

**Table S6.** Statistical significance associated with **Fig. 6** between signal-to-background (S/B) produced by various pre-miRNA-Cl sequences and pre-miR-21 with Lin28A- or Lin28B-LgBiT fusion in nuc-RiPCA (ns = not significant). Statistical significance determined by two-way ANOVA Tukey's multiple comparison test (n = 8).

| Sequence | Lin28A | Lin28B |
|---|---|---|
| **pre-miR-34a** | 0.8330 (ns) | 0.9073 (ns) |
| **pre-let-7a-1** | 0.9970 (ns) | 0.9995 (ns) |
| **pre-let-7d** | <0.0001 (****) | <0.0001 (****) |
| **pre-let-7g** | <0.0001 (****) | <0.0001 (****) |
